# The mechanical and morphological properties of systemic and pulmonary arteries differ in the earth boa, a snake without ventricular pressure separation

**DOI:** 10.1101/2021.10.07.463559

**Authors:** Benjamin J. van Soldt, Tobias Wang, Renato Filogonio, Carl Christian Danielsen

## Abstract

The walls of the mammalian aorta and pulmonary artery are characterized by diverging morphologies and mechanical properties, which has been correlated with high systemic and low pulmonary blood pressures, as a result of intraventricular pressure separation in the mammalian ventricle. However, the relation between intraventricular pressure separation and diverging aortic and pulmonary artery wall morphologies and mechanical characteristics is not understood. The snake cardiovascular system poses a unique model for the study of this question, since representatives both with and without intraventricular pressure separation exist. In this study we perform uniaxial tensile testing on vessel samples taken from the aortas and pulmonary arteries of the earth boa, *Acrantophis madagascariensis*, a species without intraventricular pressure separation. We then compare these morphological and mechanical characteristics with samples from the ball python, *Python regius*, and the yellow anaconda, *Eunectes notaeus*, species with and without intraventricular pressure separation, respectively. Strikingly, we find that although the aortas and pulmonary arteries of *A. madagascariensis* respond similarly to the same intramural blood pressures, they diverge strongly in morphology, and that this is a common attribute among species without intraventricular pressure separation in this study. In contrast, *P. regius* aortas and pulmonary arteries diverge both morphologically and in terms of their mechanical properties. Altogether our data indicate that intraventricular pressure separation does not explain diverging aortic and pulmonary artery morphologies. Following the Law of Laplace, we propose that thin pulmonary arteries represent a mechanism to protect the fragile pulmonary vascular bed by reducing the blood volume that passes through, to which genetic factors may contribute more strongly than physiological parameters.

## Introduction

Strong yet distensible arterial walls are critical for proper function of the vascular tree in animals. Strength is required to withstand high pressures when blood is ejected from the heart during systole, and distensibility is critical to ensure that the major arteries provide capacitance and pulse-pressure-smoothing after each cardiac contraction (Shadwick, 1999). These mechanical properties derive from morphological features of the vessel walls, primarily thickness, elastin and collagen content, and the extent and mode of cross-linking and alignment of the elastin and collagen fibers (Dobrin, 1978; Wagenseil et al., 2009). These properties are established during embryological development in response to hemodynamic forces, such as wall shear stress, that continuously drive vascular remodeling (Jones et al., 2006; Reneman et al., 2006). In mammals, abrupt hemodynamic changes occur after birth, when the pulmonary and systemic circuits become fully separated (Langille, 1996). Intriguingly, reports in various mammals suggest that after birth major structural changes occur in the aorta wall as compared to the pulmonary artery, the former becoming increasingly thicker-walled and stronger than the latter (Gerrity and Cliff, 1975; Leung et al., 1977). These changes probably reflect necessary adaptations to the considerably higher systemic blood pressure as compared to the pulmonary arterial pressure. However, it remains unclear whether intramural blood pressure indeed provides the causative link to differences in arterial wall morphology.

Intraventricular pressure separation describes the ability of the heart to eject blood into the systemic and pulmonary circulations at different pressures. This ability evolved independently in mammals and archosaurs (birds and crocodiles) by establishing a full ventricular septum that divides the ventricle into left (systemic) and right (pulmonary) chambers (Hicks, 1998). In contrast, with the exception of pythons and *Varanus* lizards, non-archosaur sauropsids typically lack a complete ventricular septum, resulting in similar systolic pressures in systemic and pulmonary arteries (Jensen et al., 2014). Reptiles, therefore, represent an interesting possibility to investigate the relationship between pressure separation and arterial mechanical characteristics.

In previous studies, we analyzed the mechanical characteristics of the major arteries of ball pythons (*Python regius*), which has functional intraventricular pressure separation (Jensen et al., 2010a; van Soldt et al., 2015; Wang et al., 2002; Wang et al., 2003; Zaar et al., 2007), and the yellow anaconda (*Eunectes notaeus*) that lacks intraventricular pressure separation (Filogonio et al., 2018). These studies revealed that the aortae and pulmonary arteries of *P. regius*, as in mammals, differ in their mechanical properties, while they are more similar in *E. notaeus*. However, the similarity in mechanical properties was not mirrored in morphological features, as might have been expected. Thus, it is possible that, while intraventricular pressure separation has profound effects on morphology of the great arteries, it may not play the expected causative role. In the present study, we therefore investigate the mechanical properties of the aortae and pulmonary arteries of the earth boa (*Acrantophis madagascariensis*) and compare to other species to better understand the relation between intraventricular pressure separation and the mechanical properties of the great arteries. We first demonstrate that the earth boa lacks pressure separation and then report our findings on the mechanical characteristics of the pulmonary artery and aorta walls. As in *E. notaeus*, we find that the pulmonary arteries are remarkably resilient to high strains, despite lacking apparent strength, and that this likely relates to their surprisingly small diameter, which may negate the need for increased wall strength. We conclude that differences in morphology and mechanical properties between the aorta and pulmonary arteries may be related to a combination of genetic or developmental factors in addition to pressure separation.

## Materials and methods

### Snake specimens

Nine captive-bred earth boas, *Acrantophis madagascariensis* (Duméril & Bibron, 1844), with a body mass ranging from 244–655g (344 ± 46g, mean ± SEM, Table 1) were donated by a zoological garden and kept in accordance with §53 of Danish experimental animal welfare regulations (permit ID 2013-15-2934-00847). Snakes were fed rodents weekly, but fasted several weeks prior to euthanasia.

**Table 1:**
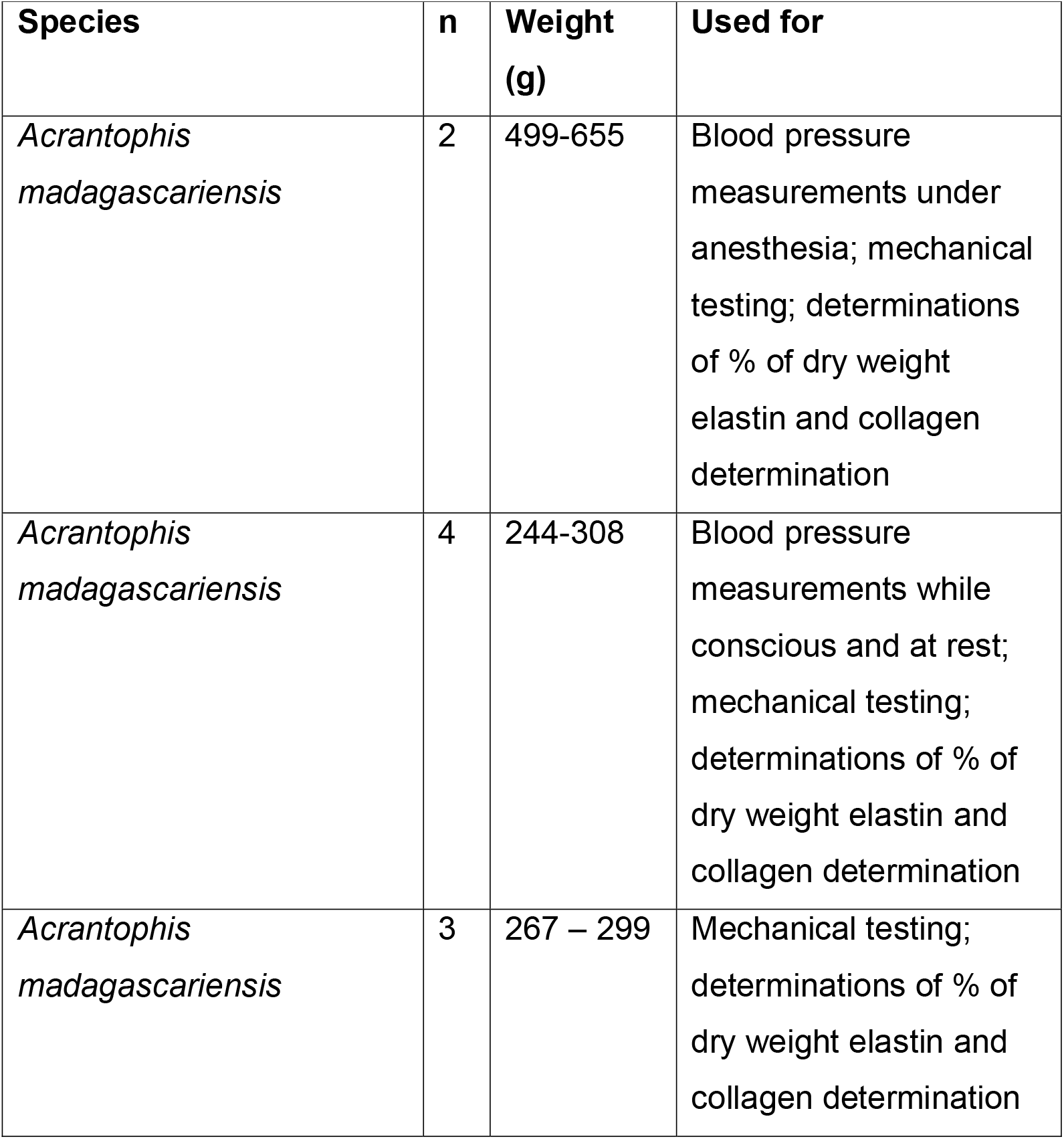
Specimens used in this study.

### Blood pressure measurements

Systemic and pulmonary blood pressures were measured in two anesthetized snakes both by intraventricular and extracardiac cannulation (vertebral and right pulmonary arteries). Snakes were anesthetized by intramuscular injection of pentobarbital (30 mg/kg) and anesthesia was confirmed by lack of muscle tone. After subcutaneous application of Xylocain (20 mg/ml), the cardiac region was exposed through a 10 cm ventral incision. Right aortic and right pulmonary arterial pressures were measured by cannulating the vertebral and right pulmonary artery with PE60 catheters containing heparinized saline (50 IU/ml). Intraventricular blood pressures were measured in the cavum arteriosum and cavum pulmonale by creating a small incision in the respective ventricular walls and inserting PE90 catheters (see Wang et al. (2003) for details on experimental procedures). To measure systemic blood pressure in four fully recovered snakes, snakes were anesthetized by inhalation of isoflurane, and the dorsal aorta was cannulated in the tail with a PE60 catheter. Pressure measurements were taken 3, 5 and 24 h after recovery from anesthesia.

Catheters were connected to Baxter Edward pressure transducers (model PX600, Irvine, CA, USA) placed at heart level of the snakes, and acquired using a Biopac MP100 data acquisition system (Goleta, CA, USA). Afterwards the snakes were euthanatized (30-50 mg/kg pentobarbital) and the heart, including great arteries, were excised. Aortic and pulmonary artery segments (∼1cm) were collected from seven locations (left and right pulmonary artery, proximal and distal locations of left and right aorta, and dorsal aorta; see also Fig. 1 in van Soldt et al. (2015)) and frozen at -20°C until further study. Freezing has minimal effect on the mechanical properties studied here (Adham et al., 1996; Chow and Zhang, 2011; Stemper et al., 2007).

**Figure 1:**
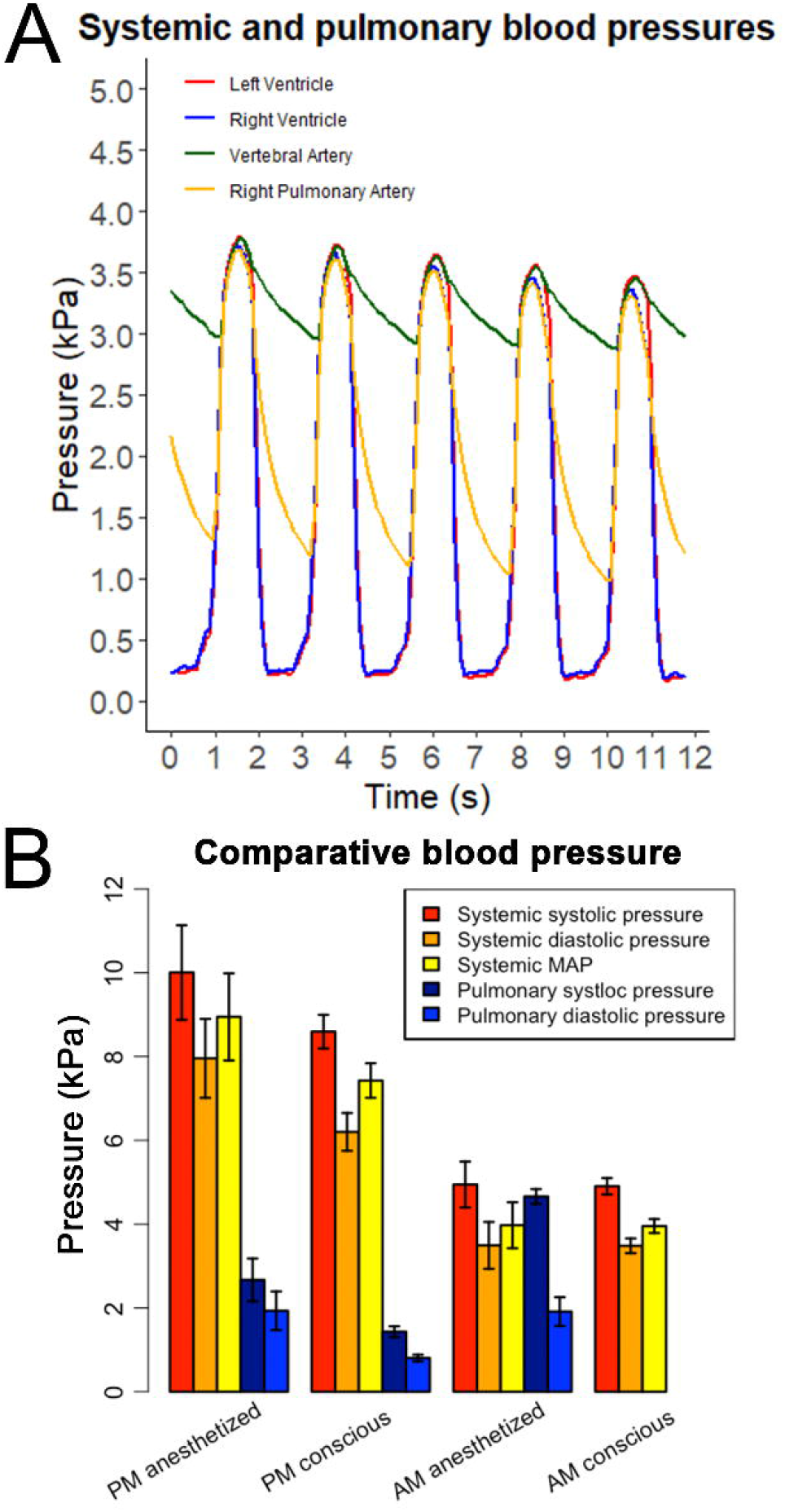
Systemic and pulmonary blood pressures in *A. madagascariensis* and *Python molurus*. (A) Waveforms showing *A. madagascariensis* left and right intraventricular blood pressures, as well as right aorta (through vertebral artery) and right pulmonary artery blood pressures and (B) bar plots comparing systemic and pulmonary systolic, mean arterial and diastolic pressures in *P. molurus* (PM) and *A. madagascariensis* (AM). Error bars in (B) indicate SEM. *P. molurus* data is from Wang et al. (2003). Note that *A. madagascariensis* was anesthetized using pentobarbital, and *P. molurus* using halothane.

### Histology

Images of histological sections of vessel segments were obtained from one snake as previously described (van Soldt et al., 2015). Briefly, the segments were fixed in formaldehyde, embedded in paraffin and sectioned (4μm). The sections were stained with resorcin, Sirius red F3B and Mayer’s haematoxylin. Photographs were taken with an Olympus C-7070 WZ camera (Tokyo, Japan) mounted on a Leica DMRB microscope (Wetzlar, Germany) using both bright field and circular polarization.

### Tissue preparation for mechanical testing

All procedures were described previously (van Soldt et al., 2015). Briefly, vessel sections were cut into rings with a nominal length of 1mm and submerged in 50mM Tris/HCl solution (pH 7.4). Cross-sectional area, height and diameter were measured by mounting the rings on a tapered glass rod at minimal strain for photography using a Nikon microscope (Tokyo, Japan) with circular polarization and analyzed using ImageJ v1.47. A scale was photographed for calibration purposes. The rings were then frozen at -20°C until further use.

### Mechanical testing

All procedures were described by van Soldt et al. (2015). Briefly, after thawing to room temperature, a vessel ring was placed around two orthogonally bent hooks with a diameter (d_h_) of 0.55mm (aortas) or 0.35mm (pulmonary arteries) and an initial linear distance between the hooks (l_h0_) of 1.2mm and 0.5mm, respectively. One hook was connected to a load cell while a step motor moved the other. Travel distance and load cell readings were acquired continuously. Each ring was subjected to a cycle of five tests with a tension maximum of 0.25N for the left or right aortas, 0.2N for the dorsal aorta, and 0.075N for pulmonary arteries, because the latter were hypothesized to rupture at lower loads. Hereafter a tension test to rupture was carried out, with maximum travel distance set at the point of vessel rupture (x). Ruptured rings were collected for hydroxyproline determination (Danielsen and Andreassen, 1988; Woessner, 1976). Collagen content was calculated as 7.46 × hydroxyproline content (Neuman and Logan, 1950). Two to four (mean 3.7) ring specimens from each vessel segment were tested.

### Determination of collagen and elastin content

To determine the elastin and collagen fractions relative to dry weight, vessels were defatted with acetone and freeze-dried. Elastin content (percentage of dry weight) was determined after an extraction procedure according to Lansing et al. (1952). Aliquots of the extracts were used for hydroxyproline determination (described above).

### Calculation of mechanical properties

Equations have been derived previously (van Soldt et al., 2015). Briefly, we first derived stress and strain values as follows:

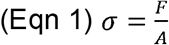

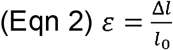

Where *F* is load, *A* is cross-sectional area, Δ*l* corresponds to incremental vessel luminal circumference (*l* = 2 · (*l*_*ho*_ +*x* − *d*_*h*_) + (*d*_*h*_ · *π*)), and *l*_0_ corresponds to vessel wall unstrained circumference, recorded as the circumference at a load value of 0.5mN. From this, we derived maximum load (*F*_max_) and strain at maximum load (*ε*_max_), as well as stress-strain and load-strain curves.

We calculated elastic modulus *E* (Gibbons and Shadwick, 1989), using the stress and strain, as follows:

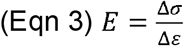

To calculate compliance curves, we first derived relative volume change (*V/V*_*0*_) from strain values. After simplification:

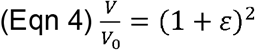

Using load (*F*) and corresponding strain (*ε*) values and the law of Laplace as applied to a cylinder we calculated pressure change (Herman, 2007), giving us:

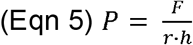

Herein, h is the nominal height, and the luminal radius (r) is 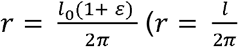 where *l*= *l*_0_(1 + *ε*)), so that:

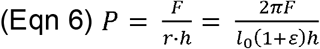

### Statistical analysis

Statistical analyses were described previously (van Soldt et al., 2015). Briefly, we ran all statistical analyses in R 4.0.3 (R Core Team, 2014) running in RStudio v1.3.1093 (RStudio, 2013), using packages lattice (Sarkar, 2008), lme4 (Bates et al., 2014), car (Fox and Weisberg, 2011) and multcomp (Hothorn et al., 2008). We used a mixed model (tested variable as dependent variable, e.g. F_max_, and individual animal (Table 1) as a random variable). A post-hoc Tukey test was used to identify significant differences between vessel segment pairs (p < 0.05). Curves were calculated for each vessel section per snake and then combined, per vessel section, into mean curves for presentation in this work. Curve data for proximal and distal segments for both aortas were pooled using described equations (Baker and Nissim 1963). One-way Anova was used for statistical tests on comparisons of morphological parameters between snake species using standard R commands. All data are displayed as mean ± SEM.

## Results

### *A. madagascariensis* lacks intraventricular pressure separation

We confirmed that *A. madagascariensis* lacks intraventricular pressure separation by measuring intraventricular and extracardiac (vertebral and right pulmonary arteries) systemic and pulmonary blood pressures (Figure 1). Systemic and pulmonary pressure waveforms overlapped entirely for intraventricular pressure measurements, both during systole and diastole. Waveforms for extracardiac measurements overlapped during systole (Figure 1A). Indeed, we observed no significant difference between systemic and pulmonary systolic pressures (4.9±0.6 and 4.7±0.2 kPa, respectively; p=0.697, n=2; Figure 1B). To ensure that anesthesia did not have significant effects on blood pressure, we also measured systemic blood pressure in conscious *A. madagascariensis*. Anesthetized (4.0±0.6 kPa, n=2) and conscious (4.0±0.2 kPa, n=4) mean arterial blood pressure (MAP) were not significantly different (p=0.978). Thus, *A. madagascariensis* does not have intraventricular pressure separation.

### The aortic walls are wider, thicker, yet more elastic than the pulmonary artery walls

We quantitively determined vessel dimensions by brightfield microscopy, combined with biochemical determinations of vessel wall composition. The aortic vessels were consistently thicker-walled (p<0.001; Figure 2A-L, M) and of larger diameter than the pulmonary arteries (p<0.0001; Figure 2A-L, N). More specifically, proximal and distal sections of left (PLAo/DLAo) and right aorta (PRAo/DRAo) were not significantly different in either wall thickness (p=1; p=0.132) or diameter (p=1; p=0.979). However, the left aorta was thicker walled (p=0.0242) and wider (p<0.0001) than the right aorta. The dorsal aorta (DAo) was of larger diameter than either the left or right aorta (p<0.00001). Finally, the right pulmonary artery (RPA) was thicker walled (p=0.006) and wider (p<0.00001) than the left pulmonary artery (LPA).

**Figure 2:**
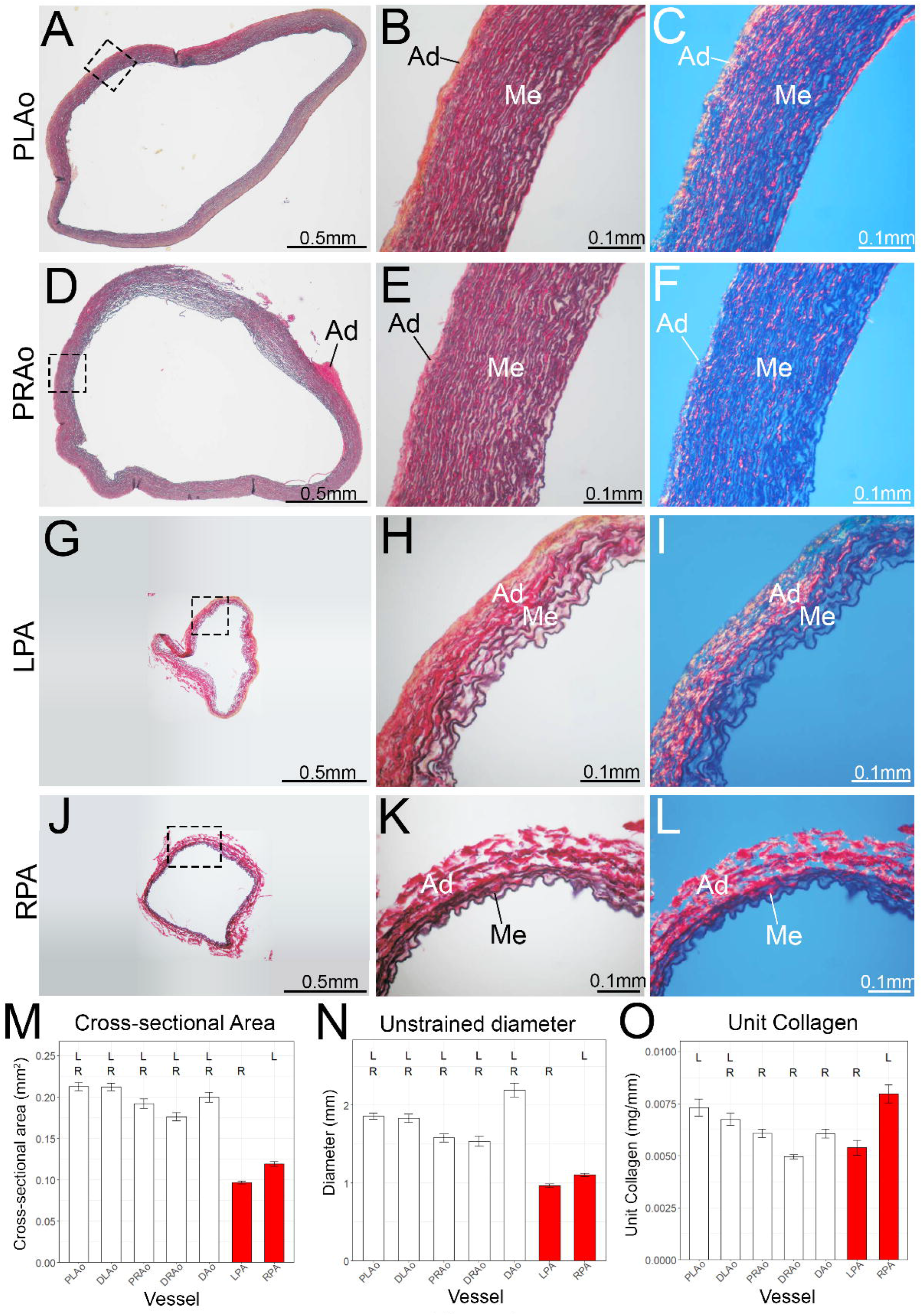
Examinations and quantifications of key histological parameters of *A. madagascariensis* aorta and pulmonary arteries. (A-L) Brightfield (A-B, D-E, G-H, J-K) and circular polarization (C,F,I,L) images of proximal sections of left (PLAo; A-C) and right aorta (PRAo; D-F), and left (LPA; G-I) and right pulmonary artery (RPA; J-L). The aortas are wider and thicker-walled than the pulmonary arteries. (M-O) Bar plots depicting vessel wall cross-sectional area (M), unstrained diameter (N) and collagen content normalized by vessel diameter (unit collagen, UC; O). “R” and “L” denote significant difference between respective vessel section and right or left pulmonary artery. Ad: tunica adventitia; Me: tunica media. Barplots display mean±SEM.

In the vessel wall, the elastin-rich tunica media confers elasticity, while the tunica adventitia—which is comprised predominantly of collagen fibers— confers strength. Microscopy using a circular polarization filter indicated that the aortic tunica media (Me; Figure 2C, F) was thicker than the tunica adventitia (Ad; Figure 2C, F). In contrast, these two layers were roughly equally thick in the pulmonary artery walls (Figure 2I, L), suggesting that the aortic walls are more elastic than those of the pulmonary arteries. Indeed, determination of elastin and collagen percentages (Table 3) showed that aortic sections had consistently higher elastin content (∼30%), but lower collagen content (∼20%) compared to pulmonary sections (∼9% and ∼40%, respectively). However, unit collagen (UC; Fig. 2O), which normalizes absolute collagen content (in mg) by vessel circumference (in mm), demonstrated that differences in absolute collagen content between aortic and pulmonary artery sections could be explained by the smaller size of the pulmonary arteries, and were consistent with identified differences in wall thickness and diameter (Figs 2M, N). Thus, the left aorta had higher UC than the right aorta (p=0.031), and the right pulmonary artery had higher UC than the left pulmonary artery (p<0.001). Altogether, these data suggest that the aortic walls are more elastic than the pulmonary artery walls.

**Table 2:** Pressure measurements for anesthetized and conscious earth boa (AM) and ball Python (PM). Measurements given as mean±SEM.

|  | Systemic (kPA) |  |  | Pulmonary (kPA) |  |  |
| --- | --- | --- | --- | --- | --- | --- |
|  | Systolic | Diastolic | MAP | Systolic | Diastolic | MAP |
| AM;<br>anesthetized | 4.94±0.5<br>5 | 3.49±0.56 | 3.97±<br>0.55 | 4.66±0.18 | 1.91±0.34 | 2.82±<br>0.74 |
| AM;<br>conscious | 4.9±0.2 | 3.48±0.18 | 3.95±<br>0.17 | N/A | N/A | N/A |
| PM;<br>anesthetized | 10±1.13 | 7.98±0.94 | 8.94±<br>1.04 | 2.67±0.51 | 1.93±0.46 | 2.26±<br>0.48 |
| PM;<br>conscious | 8.59±0.4 | 6.2±0.45 | 7.42±<br>0.41 | 1.43±0.13 | 0.8±0.08 | 1.14±<br>0.11 |

**Table 3:** Vessel ring wall composition. Samples from snakes were pooled (column 3) to return measurements per vessel section. Four determinations were performed per measurement (column 2). Elastin and collagen content as a percentage of dry weight is given as mean±SEM, when applicable.

| Vessel segment | n | Pooled samples | Elastin content (% of dry weight) | Collagen content (% of dry weight) |
| --- | --- | --- | --- | --- |
| PLAo | 2 | 1-4, 5-9 | 30.5±0.1 | 21.7±2.2 |
| DLAo | 2 | 1-5, 6-9 | 32.5±0.3 | 19.9±0.1 |
| PRAo | 2 | 1-4, 5-9 | 28.5±0.3 | 21.7±0.5 |
| DRAo | 2 | 1-4, 5-9 | 26.9±0.8 | 21.5±1.7 |
| DAo | 2 | 1-4, 5-9 | 31.0±0.0 | 17.4±0.9 |
| LPA | 1 | 1-9 | 8.9 | 39.4 |
| RPA | 1 | 1-9 | 11.1 | 41.5 |

### The aortic walls are stronger and more compliant than the pulmonary artery walls

We next investigated the mechanical properties of the aorta and pulmonary artery walls by subjecting vessel sections to 5 cycles of uniaxial mechanical tension testing, followed by a test to rupture. Loop curves demonstrated that steady state was reached by the fourth cycle for both aortic and pulmonary artery vessel segments (Figure 3A). Although load limits in cycle tests were set according to the expected maximal load that the vessel segments could endure without premature rupture (see methods), calculation of mechanical hysteresis (viscous damping, loss of elastic energy; loop area divided by area under the loading curve) showed that the pulmonary artery walls were significantly less elastic than those of the aorta (p<0.0001; Figure 3B). Pulmonary arteries also experienced higher load and stress than aortic segments at low strains, and ruptured at significantly lower load and stress values, indicating that the pulmonary artery walls are weaker than the aortic walls (p<0.001; Figure 3C-E). All aortic segments performed similarly. The left and right pulmonary artery segments also performed similarly initially, but at higher strains (ε > 0.4) the right pulmonary artery displayed steeper load/strain and stress/strain relationships than the left pulmonary artery (Figure 3D). Because left and right aortic distal and proximal sections did not differ in their mechanical properties (p=0.152, p=0.688, respectively), we pooled these sections and treated them as complete left and right aorta hereafter.

**Figure 3:**
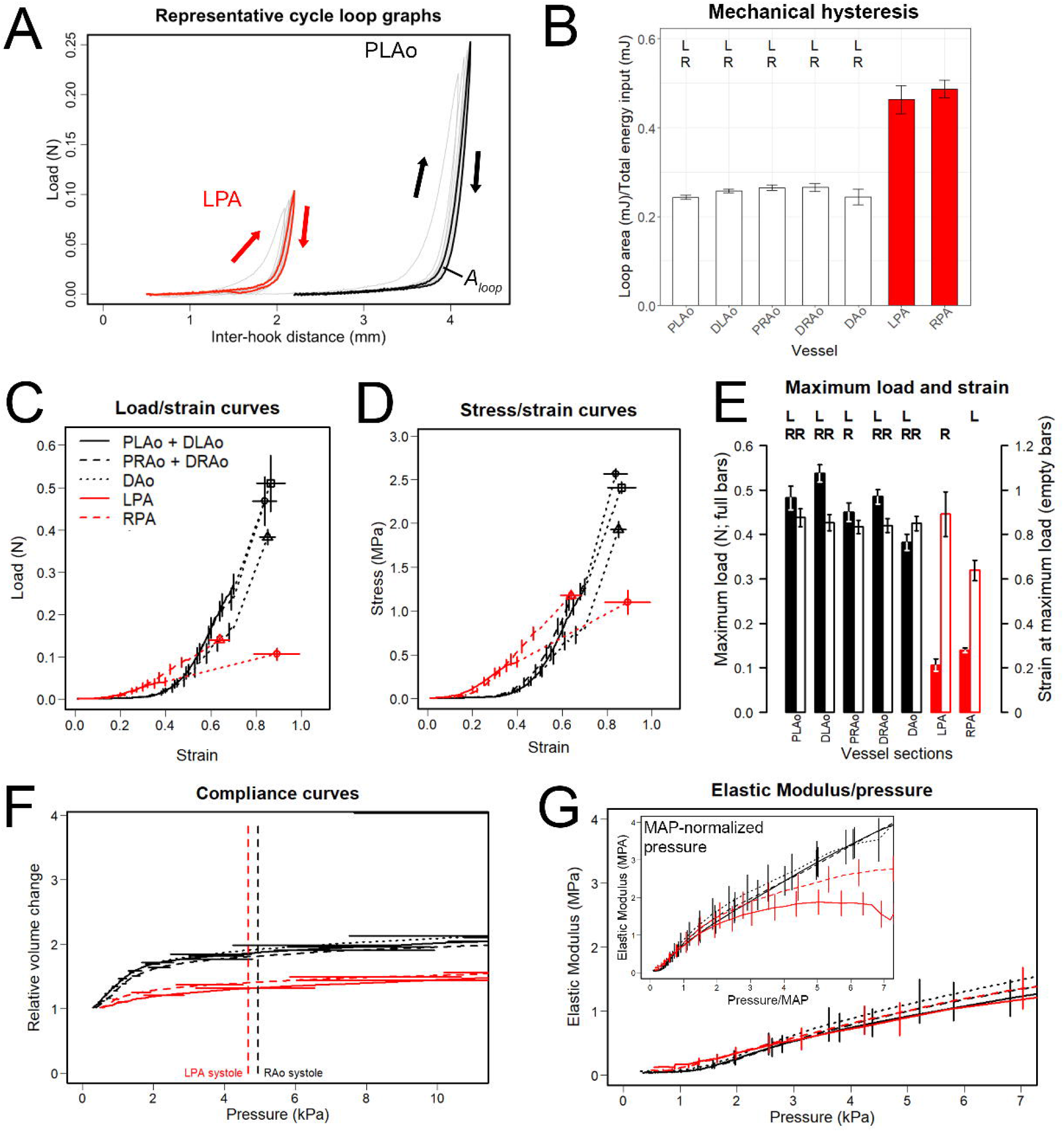
The pulmonary artery walls are weaker and less distensible than the aortic walls. (A) Representative aortic and pulmonary artery loop curves derived from 5-cycle uniaxial mechanical tension testing. Cycles one through four are grey, fifth cycle is bolded black (proximal segment of left aorta) or red (left pulmonary artery). Up and down arrows denote loading and de-loading segments of the cycle graphs, respectively. Area within loading/unloading curves is termed ‘loop area’ (*A*_*loop*_). (B) Barplot showing hysteresis (viscous damping), calculated from fifth tension test cycle (A, bolded lines) by dividing loop area (*A*_*loop*_, A) by the area under the loading curve. Load/strain curves (C), stress/strain curves (D) and barplot showing maximum load (filled bars) and strain (empty bars) (E), as well as compliance (F) and elastic modulus/pressure curves (G) derived from uniaxial mechanical tension testing to rupture. Load/strain and stress/strain curves (C) are connected by dotted lines to maximum load/strain and stress/strain values (symbols in C, D). Compliance curves include indications of systolic pulmonary and aortic blood pressures (red and black dashed lines, respectively). The elastic modulus was calculated from differentiated load/strain data divided by vessel wall cross-sectional area and plotted against pressure change (E) or pressure change normalized for mean arterial blood pressure values (E inset; 4.94kPA aortic MAP; 3.49kPA pulmonary MAP). Left and right aorta curves represent pooled data from respective proximal and distal segments. “R” and “L” denote significant difference between respective vessel section and right or left pulmonary artery. Barplots display mean±SEM.

The load a vessel wall can endure is linked to gross morphological properties, such as wall thickness and collagen content. We therefore normalized maximum load values for these two variables to obtain maximum stress and maximum load/unit collagen, the latter of which may be regarded as a measure of ‘collagen quality’ (Figure 4). In both cases, significant relationships between pulmonary artery and aorta sections persisted (p<0.001), suggesting that additional morphological factors, besides those investigated here, contribute to the differences in load that these vessels can endure.

**Figure 4:**
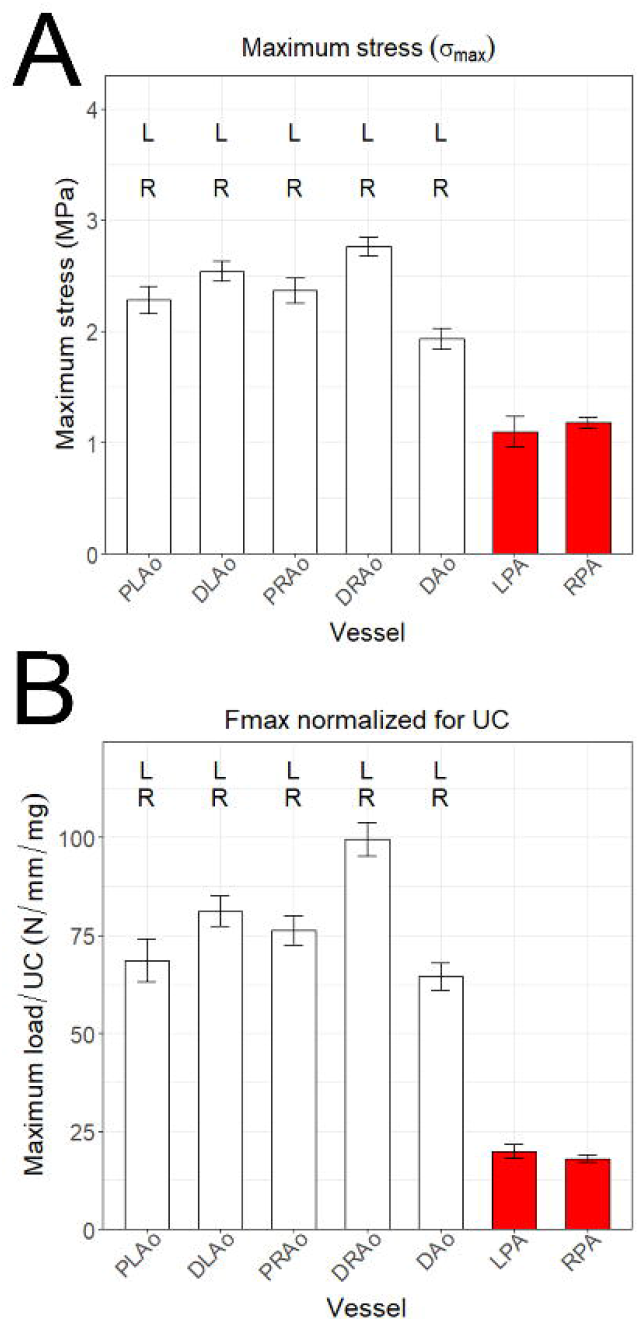
The pulmonary arteries are structurally weaker than the aortas. Barplots depict normalized values of maximum load (F_max_) for cross-sectional area (maximum stress, o_max_) (A) and unit collagen (B). “R” and “L” denote significant difference between respective vessel section and right or left pulmonary artery. Barplots denote mean±SEM.

Compliance curves calculated from load/strain curves corroborated the relative elasticity of the aortic sections (Figure 3F). While both aorta and pulmonary artery walls demonstrated a strong initial relative volume change, these were smaller for the pulmonary arteries (Figure 3F). Surprisingly, as demonstrated by plotting systolic pressures with the compliance curves, all vessels appeared to operate within the shallow portion of the curve (Figure 3F), affording them less flexibility in coping with increased blood pressure. Despite these differences in compliance, the elastic moduli, which quantify the resistance of a vessel wall to deformation, were similar for all tested vessel segments over a broad range of pressures (Figure 3G), although MAP-normalization of pressure indicated that at higher values the elastic moduli of the pulmonary arteries deviate from those of the aortas. Altogether, mechanical tensile testing of aortic and pulmonary artery segments indicated that the pulmonary artery walls were weaker and less distensible than the aortic walls, which may relate to structural features other than those investigated here.

### Intraventricular pressure separation polarizes aorta and pulmonary artery vessel wall behaviors in response to intramural blood pressures

Our data indicate that, despite the lack of intraventricular pressure separation, *A. madagascariensis* pulmonary artery and aorta walls are morphologically different, and this is reflected in the mechanical properties of these vessels. To better understand the role of intraventricular pressure separation in defining aorta and pulmonary artery mechanical characteristics, we assessed comparative vessel mechanical function in context of cardiovascular physiology of species with (*P. regius*; van Soldt et al., 2015) and without (*A. madagascariensis*) intraventricular pressure separation. We calculated vessel stiffness and strain at increasing intramural blood pressures of 2, 5 and 10kPa (Figure 5), coinciding with systemic and pulmonary systolic pressures of *A. madagascariensis* (both 5kPa) and *P. regius* (10kPa and 2.7kPa respectively; Figure 1B). Vessel strain captures deformation, while vessel stiffness describes resistance against deformation, in a manner where high stiffness correlates with low strain and vice versa.

**Figure 5:**
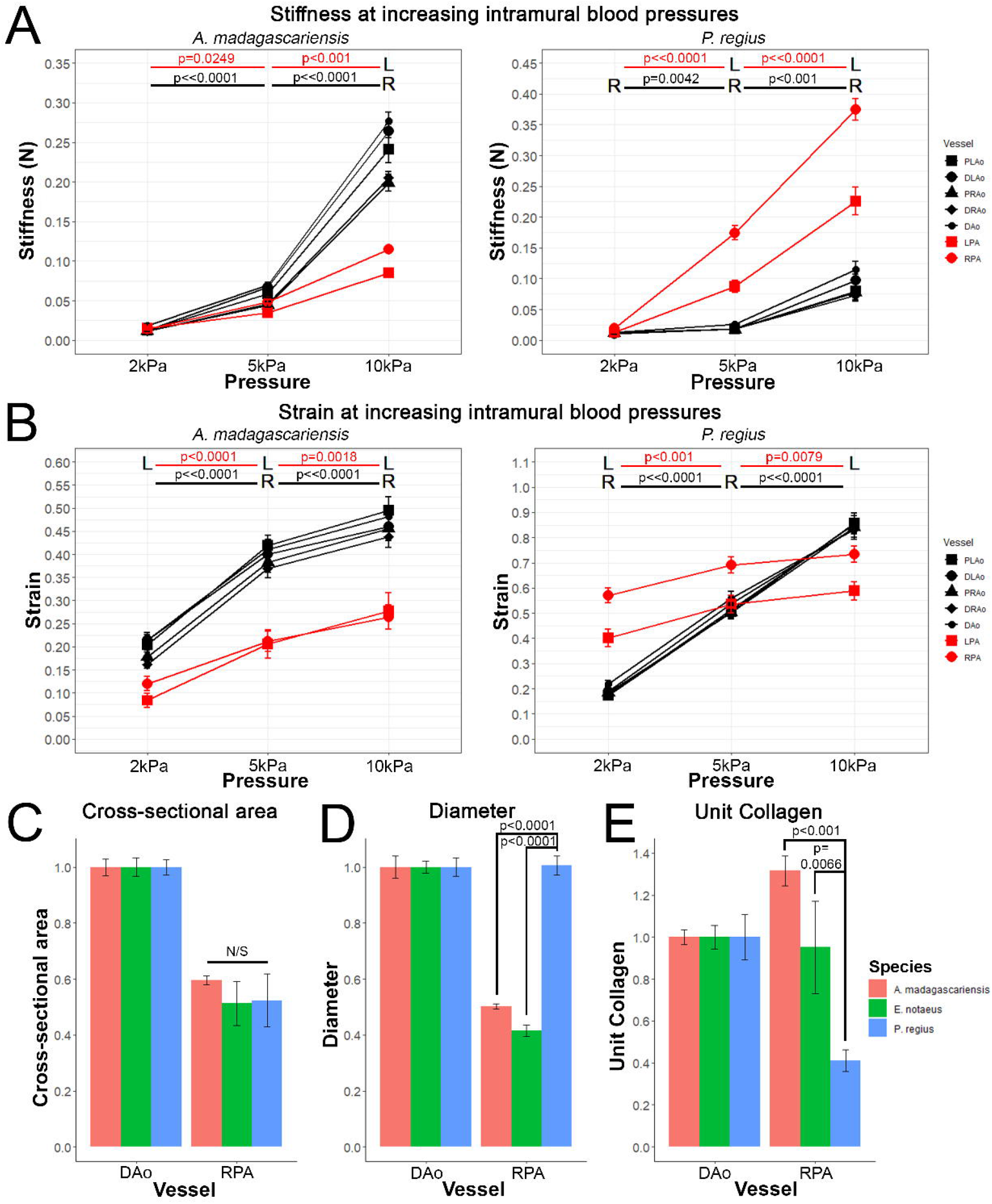
**Species comparisons of functional and morphological variables demonstrate consistent morphological and operational differences between species with and without intraventricular pressure separation**. (A-B) Line plots showing change in stiffness (A) and strain (B) of aortic and pulmonary vessels at increasing blood pressures in *A. madagascariensis* (left panels) and *P. regius* (right panels). (C-E) Barplots showing species comparisons of dorsal aorta (DAo) and right pulmonary artery (RPA) cross-sectional area (C), diameter (D) and unit collagen (E) normalized for corresponding values of dorsal aorta between *A. madagascariensis* (red bars, this study), *E. notaeus* (green bars, Filogonio et al., 2018) and *P. regius* (blue bars, van Soldt et al., 2015). “R” and “L” denote significant difference between right or left pulmonary artery and the aortic vessel sections at the corresponding intramural blood pressure, while red and black horizontal bars and p-values correspond to significant differences of stiffness and strain values between the respective intramural blood pressures (A-B). N/S denotes ‘not significant’. Data displayed as mean±SEM.

Our calculations showed that with increasing intramural blood pressure, stiffness and strain values significantly increased for all vessels regardless of species (e 5A-B). However, the relative increases in stiffness and strain from 2-5kPa and 5-10kPa differed between species and between vessels. In *A. madagascariensis* stiffness increased more steeply in aortic segments (from ∼0.025 to ∼0.06 and ∼0.235, p<0.0001; e 5A) compared to pulmonary segments (from ∼0.025 to 0.04 and ∼0.1, p=0.0249 and p<0.001, respectively; e 5A). Intriguingly, the reverse was true for *P. regius*, where stiffness increased more steeply in the two pulmonary arteries, though at different rates (on average from ∼0.025 to ∼0.15 and ∼0.3, p<0.0001; e 5A), compared to the aortas (∼0.025 to ∼0.028 and ∼0.09. p=0.0042 and p<0.001, respectively; e 5A). In regard to strain, *A. madagascariensis* aortas and pulmonary arteries appeared to operate at significantly different levels of strain but strain increased similarly in response to pressure increases (∼2.5 times from 2kPa to 10kPa, p<0.0001; e 5B). In *P. regius*, however, the pulmonary arteries operate at higher strains than the aortas at 2kPa (p<0.001; e 5B) and only display, on average, a ∼20% increase in strain at 10kPa. In addition, the relative increase in strain is comparatively steeper from 2-5kPa (p<0.001; e 5B) than from 5-10kPa (p=0.0079; e 5B). In contrast, the aortas display significant, linear increases in strain from 2-10kPa (p<0.0001; e 5B). Combined, these data show that in both species the increase in vessel stiffness is steeper when blood pressure passes above species-specific systolic pressures, resulting in a polarization of vessel mechanical behaviors in *P. regius*, where systemic and pulmonary blood pressures are significantly different.

We hypothesized that the observed inter-species differences were associated with differences in vessel morphologies. We therefore compared morphological parameters (cross-sectional area, diameter and unit collagen) of vessel walls between these species (Figure 5C-E). Intriguingly, while wall cross-sectional area did not differ (Figure 5C), *A. madagascariensis* had significantly narrower pulmonary arteries (p<0.0001; Figure 5D) with a higher unit collagen (p<0.01; Figure 5E) than *P. regius*.

Thus, *A. madagascariensis* pulmonary arteries are significantly narrower and weaker than aortas, but nevertheless respond similarly to increases in blood pressure. This stands in contrast to *P. regius*, where aortas and pulmonary arteries respond differently to increases in blood pressure.

## Discussion

Thoma was first to propose that blood circulation impacts blood vessel morphogenesis, describing relationships between blood flow and vessel radius, as well as blood pressure and wall cross-sectional area (Thoma, 1893; Wagenseil and Mecham, 2009). Indeed, the wall of the mammalian aorta is thick and strong in comparison to the pulmonary artery, and these differences are associated with high systemic and low pulmonary blood pressures as a result of intraventricular pressure separation postnatally (Gerrity and Cliff, 1975; Leung et al., 1977). We here showed measurements from *Acrantophis madagascariensis* that demonstrate that morphological and mechanical characteristics of aorta and pulmonary artery can diverge even in absence of intraventricular pressure separation, corroborated by data from *E. notaeus* (Filogonio et al., 2018).

The absence of intraventricular pressure separation in *A. madagascariensis* is obvious when considering data from *Python molurus* (Figure 1B; Wang et al., 2003). Here, systemic systolic pressure (10.01±1.13 kPa) is higher than pulmonary systolic pressure (2.67±0.51 kPa). Importantly, *P. molurus* systemic systolic pressure was higher (p=0.032) and pulmonary systolic pressure lower (p=0.049) than in *A. madagascariensis*.

Despite differences in morphology and mechanical characteristics between *A. madagascariensis* aorta and pulmonary artery walls, these vessels nevertheless responded similarly to increasing wall tension within physiological blood pressure ranges. In contrast, *P. regius* aorta and pulmonary artery responded differently, consistent with the intraventricular pressure separation in this species that polarizes systemic and pulmonary blood pressures. For example, the stiffness of all *A. madagascariensis* vessels increased more steeply from 5-10kPa than 2-5kPa, in line with a blood pressure of 5kPa (Figure 5A,B). In contrast, the P. regius pulmonary artery stiffness increased dramatically at pressures over 2kPa, reflecting the systolic pulmonary blood pressure of 2.7kPa. This aligns with stiffness/strain calculations in human and pig, where the pulmonary artery was also stiffer than the aorta at systemic blood pressures (Azadani et al., 2012; Matthews et al., 2010). Thus, the aorta and pulmonary artery are each optimized to handle respective physiological blood pressures, regardless of whether these are equal or divergent.

We were initially surprised that *A. madagascariensis* pulmonary arteries withstood the same blood pressures as the aorta, given their lower strength. However, the law of Laplace intuitively explains that narrow pulmonary arteries do not require strong, thick walls (high cross-sectional area) (Burton, 1965; Shadwick, 1999; Valentinuzzi and Kohen, 2011). Indeed, the pulmonary arteries of *A. madagascariensis* and *E. notaeus* were narrower than in *P. regius*, but with similar cross-sectional areas (Figure 5C-D). Thus, the reduction of pulmonary artery radius may be a mechanism to withstand high pulmonary blood pressure in species that lack intraventricular pressure separation.

While theoretically compelling, the narrow pulmonary arteries of *A. madagascariensis* and *E. notaeus* were surprising, since narrow vessels implicate low blood volume. Studies in sheep demonstrated that abdominal blood flow and vessel diameter decreased concurrently postnatally (Bendeck and Langille, 1992; Bendeck et al., 1994; Langille, 1996; Langille et al., 1990). In rabbit, aorta and pulmonary artery diameters remain similar postnatally while aorta wall thickness increases, likely to accommodate increasing aortic blood pressure (Leung et al., 1977). Likewise, in *P. regius* we previously found that aorta and pulmonary artery diameters were similar, but the aortic wall was thicker (van Soldt et al., 2015). Thus, one possible explanation for the narrow pulmonary arteries in *A. madagascariensis* and *E. notaeus* includes the capacity for right-to-left cardiac shunts, whereby systemic blood bypasses the lungs for systemic recirculation, decreasing pulmonary blood flow (Hicks, 1998). *P. regius* has limited capacity for such shunts (Jensen and Wang, 2009; Jensen et al., 2010b; Wang et al., 2003), and in mammals such shunts are impossible due to physical separation of systemic and pulmonary circuits (Hicks, 1998). Thus, these animals may require a wider pulmonary artery to accommodate the volume of blood flowing through these vessels, instead gaining a thicker aortic vessel wall to accommodate the higher systemic blood pressure. Indeed, In the American alligator, *Alligator mississipiensis*, a species with complete ventricular separation but with an ability to promote right-to-left shunts through the foramen of Panizza, the left pulmonary artery is narrower than the right aorta, corroborating the hypothesis (Filogonio et al., 2021). However, further study is required to ascertain the level of intracardiac shunting in *A. madagascariensis* and *E. notaeus*.

We showed that pulmonary artery unit collagen was markedly higher in *A. madagascariensis* and *E. notaeus* compared to *P. regius*. Blood vessel wall structure is fundamentally similar across species, but collagen and elastin content are known to differ to change vessel mechanical properties (Shadwick, 1998; Shadwick, 1999). Wall strength is primarily mediated by collagen, suggesting that high unit collagen may complement reduced vessel diameter to strengthen the pulmonary arteries (Dobrin, 1978; Sage and Gray, 1979). Structural features of collagen that were not analyzed here, such as cross-linking and fiber alignment, could further contribute to a role for collagen in strengthening pulmonary artery walls. Other components of the extracellular matrix, such as glycosaminoglygans, may also be determinant in defining several morphological and mechanical characteristics of the arterial wall (Gandley et al., 1997).

Given that intraventricular pressure separation alone may not define gross morphological characteristics of the aorta and pulmonary artery, we suggest that ontogenetic factors may also contribute. Importantly, the aorta and pulmonary arteries derive from different cellular progenitor populations (DeSesso, 2017; Herriges and Morrisey, 2014; Peng and Morrisey, 2013; Peng et al., 2013). Thus, this divergent ontogeny may lay the foundation for diverging transcriptional programs that are ultimately expressed in the different morphology of the pulmonary artery as compared to the aorta.

In conclusion, we showed that the absence of intraventricular pressure separation in *A. madagascariensis* does not equalize the morphologies and mechanical characteristics of its aorta and pulmonary artery. We propose that, to mitigate a higher wall tension as a consequence of increased pulmonary blood pressure, the pulmonary artery became narrower and the collagen content of its wall increased. However, this may only be possible in species with a capacity for right-to-left cardiac shunts as a mechanism to decrease pulmonary blood flow. Finally, we suggest that ontogenetic differences may be fundamental to the morphological differences between these arteries. Thus, in evolution, compensatory mechanisms to accommodate a range of intramural blood pressures may have followed the law of Laplace in different and sometimes unexpected ways.

## List of Symbols and Abbreviations

Ad: tunica adventitia
d_h_: hook diameter
DAo: dorsal aorta
DLAo: distal section of the left aorta
DRAo: distal secion of the right aorta
l_h0_: linear distance between hooks
LPA: left pulmonary artery
MAP: mean arterial blood pressure
Me: tunica media
PLAo: proximal section of the left aorta
PRAo: proximal section of the right aorta
RPA: right pulmonary artery
SEM: standard error of the mean
UC: unit collagen
x: hook travel distance at point of vessel rupture

## Acknowledgements

The authors declare no conflict of interest. We thank Mrs. Jytte Utoft for cutting the vessel sections for histology, Mrs. E. K. Mikkelsen for her help in the elastin and hydroxyproline determinations, and Dr. H. G. J. van Mil for his expert help with the statistics. We also thank Dr. C. Williams for measuring blood pressures in anesthetized *A. madagascariensis* specimens. The authors were supported by the Danish Council for Independent Research (Det Frie Forskningsråd|Natur og Univers), the Leiden University Fund (LUF), the Royal Dutch Zoological Society (KNDV), the Outbound Study Grant (OSG) as well as the Erasmus student exchange program.

